# Mid-Infrared Spectroscopic Analysis of Raw Milk to Predict the Blood Plasma Non-Esterified Fatty Acid Concentration in Dairy Cows

**DOI:** 10.1101/853127

**Authors:** Ben Aernouts, Ines Adriaens, José Diaz-Olivares, Wouter Saeys, Päivi Mäntysaari, Tuomo Kokkonen, Terhi Mehtiö, Sari Kajava, Paula Lidauer, Martin H. Lidauer, Matti Pastell

**Affiliations:** KU Leuven, Department of Biosystems, Biosystems Technology Cluster, Campus Geel, Kleinhoefstraat 4, 2440 Geel, Belgium; KU Leuven, Department of Biosystems, Mechatronics, Biostatistics and Sensors division, Kasteelpark Arenberg 30, 3001 Leuven, Belgium; Natural Resources Institute of Finland (Luke), Maarintie 6, 02150 Espoo, Finland; Natural Resources Institute of Finland (Luke), Tietotie 4, 31600 Jokioinen, Finland; University of Helsinki, Department of Agricultural Sciences, Koetilantie 5, 00014 Helsinki, Finland; Natural Resources Institute of Finland (Luke), Halolantie 31 A, 71750 Maaninka, Finland

**Keywords:** milk mid-infrared spectroscopy, blood plasma non-esterified fatty acid concentration, negative energy status, milk biomarker

## Abstract

In high yielding dairy cattle, severe postpartum negative energy status is often associated with metabolic and infectious disorders that negatively affect production, fertility and welfare. Mobilization of adipose tissue associated with a negative energy status is reflected through an increased level of non-esterified fatty acids (**NEFA**) in the blood plasma. Earlier, identification of a negative energy status through the detection of increased blood plasma NEFA concentration required laborious and stressful blood sampling. More recently there have been attempts to predict blood NEFA concentration from milk samples. This study aimed to develop and validate a model to predict the blood plasma NEFA concentration using milk mid-infrared (**MIR**) spectra that are routinely measured in the context of milk recording. To this end, blood plasma and milk samples were collected in weeks 2, 3 and 20 post-partum for 192 lactations in 3 different herds. The blood plasma samples were taken in the morning, while representative milk samples were collected during the morning and evening milk session on the same day. To predict the blood plasma NEFA concentration from the milk MIR spectra, partial least squares regression models were trained on part of the observations from the first herd. The models were then thoroughly validated on all other observations of the first herd and on the observations of the two independent herds to explore their robustness and wide applicability. The final model can accurately predict blood plasma NEFA concentrations below 0.6 mmol/L with a root mean square error of prediction (RMSE) of less than 0.143 mmol/L. However, for blood plasma with more than 1.2 mmol/L NEFA, the model clearly underestimates the true level. Additionally, it was found that morning blood plasma NEFA levels were predicted with a significantly higher accuracy (*p* = 0.009) using MIR spectra of evening milk samples compared to morning samples, with RMSEP values of respectively 0.182 and 0.197 mmol/L and *R*^2^ values of 0.613 and 0.502. These results suggest a time delay between variations in blood plasma NEFA and related milk biomarkers. Based on the MIR spectra of evening milk samples, cows at risk for a negative energy status, indicated with detrimental morning blood plasma NEFA levels (> 0.6 mmol/L), could be identified with a sensitivity and specificity of respectively 0.831 and 0.800. As this model can be applied to millions of historical and future milk MIR spectra, it opens opportunities for regular metabolic screening and improved resilience phenotyping.

## INTRODUCTION

The transition from pregnancy to lactation in high-yielding dairy cows is typically accompanied by a negative energy status (**ES**) in which the energy requirement exceeds the energy input from feed. As severe negative ES increases the susceptibility to various health and fertility problems (Leblanc, 2010; Ospina et al., 2010a), the duration and degree of negative ES should be limited through preventive actions in combination with individual monitoring and imperative treatment.

To compensate for the energy deficit and maintain high milk production, adipose tissue is mobilized and non-esterified fatty acids (**NEFA**) are released in the blood. Hence, a blood plasma NEFA concentration above 0.6 mmol/L is generally used as an indicator for negative ES in dairy cattle (Ospina et al., 2010b). These high concentrations of circulating NEFA have a detrimental effect on the oocyte quality and the immune response of dairy cows (Leroy et al., 2005; Scalia et al., 2006). In the liver, part of the NEFA are oxidized completely to deliver energy or incompletely to produce ketone bodies (Adewuyi et al., 2005). Another portion of the NEFA is esterified to triglycerides and either stored in the liver or transported as lipoproteins to e.g. the alveolar epithelial cells of the udder tissue to synthetize milk fat. In this way, fatty acids (**FA**) and ketone bodies derived from the NEFA end up in the produced milk. Previous studies have demonstrated the use of milk biomarkers for monitoring negative ES in individual cows, e.g. through the measurement of certain FA (Van Haelst et al., 2008; Jorjong et al., 2014; Dórea et al., 2017), ketone bodies (Enjalbert et al., 2010), citrate and many more (Bjerre-Harpøth et al., 2012). In contrast to taking blood samples, milk sampling requires less labor and can be done without distressing the animals. Nevertheless, the reference techniques to measure these milk biomarkers are typically labor-intensive and costly (Jorjong et al., 2014).

A relatively straightforward and cost-efficient technique for milk analysis is mid-infrared (**MIR**) spectroscopy. As the covalent bonds of molecules in milk absorb MIR radiation at very specific wavenumbers, the concentrations of these milk components can be derived from the MIR absorbance spectra. Typically, multivariate linear models are trained to predict the milk constituents from the acquired spectra (De Marchi et al., 2014). Already for decades, this technique is accepted as the reference for accurate and routinely characterization of the main milk components in the context of milk recording (ISO, 2013; ICAR, 2019). Since the commercial introduction of Fourier-transform MIR spectrometers for milk analysis, milk MIR spectra can be obtained with a higher accuracy and repeatability. This opens opportunities for measuring minor milk components and milk biomarkers such as FA profiles (Rutten et al., 2009; Afseth et al., 2010; Soyeurt et al., 2011), protein composition (Franzoi et al., 2019), minerals (Soyeurt et al., 2009), ketone bodies and citrate (Grelet et al., 2016).

Recently, Benedet et al. (2019), Grelet et al. (2019) and Luke et al. (2019) developed models to predict the blood plasma NEFA concentrations from milk MIR spectra of individual dairy cows. However, the prediction performance of Grelet’s model was poor (*R*^2^ = 0.39), likely because it was built using a limited number (*n* = 234) of calibration samples (Grelet et al., 2019). Benedet’s model performed better (*R*^2^ = 0.52), however, like Grelet’s model, it was not validated for a completely independent herd (Benedet et al., 2019; Grelet et al., 2019). Accordingly, the reported results might be overoptimistic compared to applying the model on the data of a new herd where the cows are managed differently. This was clearly illustrated by Luke et al. (2019) as the determination coefficient (*R*^2^) of their model dropped from 0.61 for a randomly selected validation set, covering the same herds as the ones included in the calibration set, to 0.45 for a completely independent herd.

We hypothesized that a better prediction performance can be obtained through increasing the number of calibration samples and applying a very strict timing in the sampling of blood and milk samples relative to the diurnal pattern and the feeding schedule of the cows (Quiroz-Rocha et al., 2010). To test this hypothesis, a high number of samples was collected following a strict protocol for blood and milk sample collection to obtain high quality data for training the prediction models. Additionally, it is investigated whether MIR spectra of morning or evening milk samples result in a better prediction of the NEFA concentration of the respective blood samples taken in the morning of that day. Finally, the performance of the prediction models is evaluated extensively on a completely independent validation set.

## MATERIALS AND METHODS

### Experimental Setup

The experimental protocol was approved by the Finnish Animal Experiment Board (ESAVI/5688/04.10.07/2013) and applied on 3 experimental herds in Finland: Luke Jokioinen (herd A), University of Helsinki in Viikki (herd B) and Luke Kuopio (herd C). All cows in these herds that calved for the first time in the period between September 2013 and October 2016 were included in the study, resulting in a total of 143 Nordic Red dairy cows from which 103 were in herd A, 24 in herd B and 16 in herd C. For 49 of these 143 cows, also the second lactation was included in the study period, thus resulting in a total of 192 lactations. A detailed description on the housing conditions, ration, feeding frequency and milking conditions and frequency during the experiment is given by Mäntysaari et al. (2019).

### Data Collection

For each lactation, blood samples were taken from the coccygeal vein within one hour after the morning milking session, on two non-consecutive days in week 2 and in week 3 after calving, and once in week 20. This resulted in a total of 5 blood samples per lactation. Handling of the lactating cows prior to blood-sampling was minimized to reduce its effect on the blood plasma NEFA concentrations (Leroy et al., 2011). Blood was collected in 10 mL EDTA tubes and stored in ice until centrifuged at −4°C for 15 min at 2,000 × *g*. Plasma samples were frozen and stored at −20°C for later analysis of NEFA at the university of Helsinki (Salin et al., 2012). An enzymatic colorimetric acyl-CoA synthetase (ACS)-acyl-CoA oxidase (ACOD) method [NEFA-HR(2) kit, Wako Chemicals GmbH, Neuss, Germany] was used according to the manufacturer’s instructions to determine the blood plasma NEFA concentrations, further referred to as ‘blood NEFA’. Intra- and interassay coefficient of variation for blood NEFA determination were 1.61 and 3.53% for low NEFA concentration (0.23 mmol/L) and 0.77 and 2.91% for high NEFA concentration (1.24 mmol/L).

Representative milk samples (± 30 mL) were collected during the morning and evening milking sessions on the same days as the blood collection, providing a total of 10 milk samples per lactation. The milk samples were stored at 4°C using a preservative (± 0.3 mg bronopol per ml milk, Broad Spectrum Microtabs II, D and F Control Systems Inc., Dublin, CA). The MIR analyses (MilkoScan FT6000 spectrometer, Foss, Hillerød, Denmark) were carried out by the Valio Ltd. milk laboratory (Seinäjoki, Finland) according to ISO 9622:2013 (ISO, 2013). The MIR spectrum of each milk sample consisted of 1060 values, representing the infrared light transmittance through 50 µm of sample between wavenumbers 5010.2 and 925.7 cm^−1^ with a resolution of 4 cm^−1^. The MIR spectra were standardized following the procedure developed by Grelet et al. (2015). Because of data storage problems, the MIR spectra of 152 morning milk samples and 183 evening milk samples got lost. The resulting final dataset therefore included 808 and 777 MIR spectra for respectively morning and evening milk samples (Table 1).

**Table 1.** Number of mid-infrared (MIR) transmittance spectra of morning and evening milk samples available for the 3 different herds included in this study.

| Milk samples | All herds | Herd |  |  |
| --- | --- | --- | --- | --- |
|  |  | A | B | C |
| Morning | 808 | 640 | 121 | 47 |
| Evening | 777 | 646 | 118 | 13 |

### Prediction of the Blood Plasma NEFA Concentrations from Milk MIR Spectra

The MIR spectra and NEFA concentrations were imported into R version 3.4.3 (R Core Team, 2017). Only the spectral regions from 2977 to 2768 cm^−1^, 1800 to 1684 cm^−1^ and 1607 to 926 cm^−1^ were used in the analysis. Moreover, the signal-to-noise ratio in the spectral regions between 3660 and 2977 cm^−1^ and between 1684 and 1607 cm^−1^, was considered too low due to substantial MIR absorption by the water molecules. The spectral regions above 3660 cm^−1^ and between 2768 and 1800 cm^−1^ were deleted because they do not contain significant spectral information on relevant milk components (Aernouts et al., 2011; Grelet et al., 2019). A principal component analysis (**PCA**) with maximum 20 principal components was used to identify potential outlier spectra. When both the *Q* residuals and the Hotelling *T^2^* statistic were above their 99% confidence limits, the spectrum was removed from the analysis (Bro and Smilde, 2014).

As blood samples were only taken once per cow per sampling day, while 2 milk samples were collected for respectively the morning and evening milking session of that day and cow, the number of blood NEFA analyses was half of the amount of milk MIR spectra. Accordingly, the same blood NEFA concentration was assigned to both the morning and the evening milk MIR spectrum of the respective cow and day. The combination of a morning milk MIR spectrum together with the respective blood NEFA concentration is further referred to as a morning observation, while the combination of an evening milk MIR spectrum together with the respective blood NEFA concentration is further referred to as an evening observation. The morning and evening observation of the same cow and day thus have the same blood NEFA concentration, while they have different milk MIR spectra. Next, about 60% of the morning and evening observations of herd A were allocated to the calibration set, while the remaining 40% of the observations of herd A and all observations of herds B and C were assigned to the validation set (Figure 1, step 1). Moreover, the observations of herd A were split 60/40 by applying the duplex selection method after ordering them on their blood NEFA concentration (Snee, 1977). This procedure assured that both sets had similar descriptive statistics. Observations for the same cow were treated as a block with all of them either in the calibration or validation set to prevent overoptimistic validation results in case of modeling cow-specific effects (Kemps et al., 2010).

**Figure 1.**
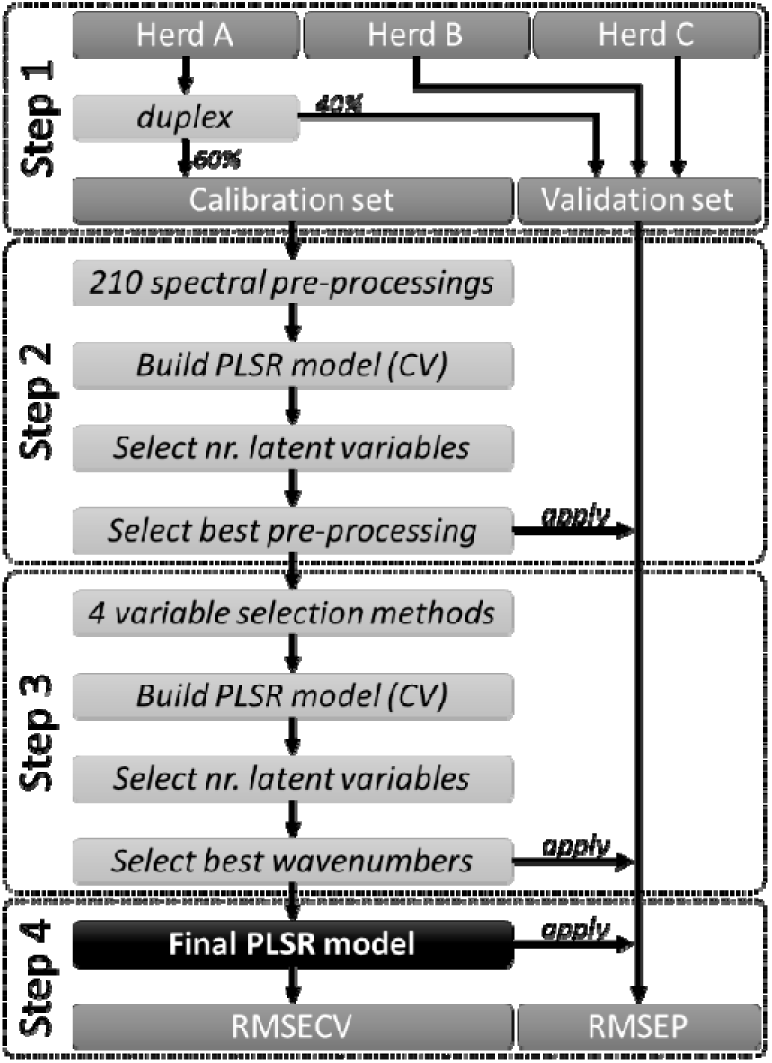
Schematic overview of the methodology to build a partial least squares (PLSR) model to predict the blood plasma non-esterified fatty acid concentration from milk mid-infrared spectra. CV = cross-validation, RMSECV = root mean square error of cross-validation, RMSEP = root mean square error of prediction.

The spectral pre-processing of the MIR spectra was a combination of (1) a logarithmic spectral transformation (Beer, 1852) or not; (2) a baseline correction, detrending, standard normal variates weighting or multiplicative scatter correction (Geladi et al., 1985; Barnes et al., 1989; Ruckstuhl et al., 2001) or none of those; (3) a first or second order Savitzky-Golay derivative with a second order polynomial filter and 10 different spectral window lengths (Savitzky and Golay, 1964) or no derivative and (4) mean centering. This resulted in 210 different combinations, as presented in Figure 1 (step 2) and described in detail in Aernouts et al. (2011). For each of these 210 combinations, a partial least squares regression (**PLSR**) model with up to 20 latent variables was built to predict the blood plasma NEFA concentrations, further referred to as ‘predicted blood NEFA’, from the pre-processed MIR spectra (Martens and Næs, 1987). A group-wise cross-validation (**CV**) with 20 groups, each containing spectra of 3 to 4 cows, was performed on the observations of the calibration set to obtain the root mean square error of cross-validation (**RMSECV**). We selected the smallest number of latent variables for which the PLSR model was not significantly worse compared to the same model with the number of latent variables resulting in the lowest RMSECV. The statistical comparison in this procedure was based on a one-sided paired *T*-test (α = 0.05) applied on the absolute residuals of the cross-validated observations (Cederkvist et al., 2005). A similar approach was followed to select the best spectral pre-processing combination. Moreover, the PLSR models resulting from the 210 combinations were ranked by increasing RMSECV, and the one with the smallest number of latent variables and not being significantly worse compared to the model with the lowest RMSECV was selected. Again, a one-sided paired *T*-test (α = 0.05) on the absolute residuals of the cross-validated observations was used to statistically compare the models (Cederkvist et al., 2005; Aernouts et al., 2011).

The selected pre-processing combination was applied on the MIR spectra to be used as an input for 4 different variable selection methods (Figure 1, step 3): variable importance in projection, jack-knife, reversed interval PLSR and forward interval PLSR (Norgaard et al., 2000; Westad and Martens, 2000; Chong and Jun, 2005). Each of these 4 methods resulted in a set of most relevant wavenumbers for which a PLSR model with an optimal number of latent variables was built as described earlier. The performances of these 4 PLSR models were compared mutually and with the model that uses all wavenumbers. Finally, the set of wavenumbers related to the most parsimonious model whose prediction performance was not significantly worse (one-sided paired *T*-test, α = 0.05) than that of the model with the lowest RMSECV was selected.

The final prediction model (Figure 1, step 4), together with the selected combination of spectral pre-processing techniques and the selected set of wavenumbers, was used to predict the NEFA concentrations of the observations in the validation set. Accordingly, an error or residual could be calculated for each observation of the validation set. Based on these residuals, the root mean square error of prediction (**RMSEP**), further referred to as the ‘prediction error’, was calculated for the entire validation set. Because this validation set is very diverse, containing morning and evening observations from 3 different herds with blood NEFA concentrations ranging from very low to very high, the RMSEP was also calculated for different subsets of the validation set, allowing for a better understanding of the prediction performance of the model under different situations. These subsets were defined based on a combination of the following features:

- Milking time: only morning observations, only evening observations or both morning and evening observations;
- Herd: observations from herd A, herd B, herd C or for the 3 herds together;
- NEFA range: observations with blood NEFA concentrations in the low (0 – 0.6 mmol/L), middle (0.6 – 1.2 mmol/L), high (1.2 – 2.0 mmol/L) or complete range (0 – 2 mmol/L). These ranges were defined like this because 0.6 mmol/L is generally considered as critical threshold (Ospina et al., 2010b) and because the blood NEFA concentration was always underestimated for true concentrations above 1.2 mmol/L.

The procedure described above (Figure 1) was initially followed to develop and validate a PLSR model that predicts the blood NEFA independent of the moment of milk sampling by training it on all the observations – both morning and evening – of the calibration set. This model is further referred to as the ‘full model’. To evaluate the effect of restricting the calibration set to only morning or evening observations, 2 new models were trained following the same procedure as elaborated above, but with respectively only the morning or the evening observations of the calibration set for training the respective PLSR models. These models are further referred to as respectively the ‘morning model’ and the ‘evening model’. All 3 models (full, morning and evening) were validated on the same observations – both morning and evening – of the validation set to allow for an objective comparison of the prediction performance.

The prediction performances of the 3 models were compared by applying a repeated-measures ANOVA on the absolute residuals for all the observations of the validation set. Moreover, ‘model’ was treated as a fixed effect, while ‘sample’ was specified as a random effect in the two-way ANOVA (Cederkvist et al., 2005). When the ANOVA test pointed out a significant effect (α = 0.05) of the model, then the performance of the 3 models was compared bilateral using a Tukey HSD multiple comparison (α = 0.05). The 3 models were compared for all the observations in the validation set, as well as the observations in the different subsets of the validation set. The model (full, morning or evening) which was not significantly different from the best model for most of the subsets of the validation set was identified as the most robust. This model was further evaluated on its ability to identify detrimental blood plasma NEFA concentrations (next section). Finally, a 4-way ANOVA analysis, with the model, the milking time, the herd, the NEFA range and all possible interactions as fixed factors, was applied on the absolute residuals for the observations of the entire validation set and subsets of the validation set. This analysis was not paired, so the samples could not be taken as a random factor. If one of the interactions was significant (α = 0.05) then all possible combinations of the factors involved in these interactions were compared bilateral using the Tukey HSD multiple comparisons. In absence of significant interaction for a factor, the effect of the factors could be interpreted separately. Moreover, if this factor had a significant (α = 0.05) influence on the performance, then the different levels within this factor were compared bilateral with the Tukey HSD multiple comparisons.

### Identify Detrimental Blood Plasma NEFA Concentrations from Milk MIR Spectra

To evaluate whether the predicted blood NEFA concentrations can be used to identify detrimental blood NEFA levels (> 0.6 mmol/L), receiver operating characteristic (**ROC**) analyses were performed (Ospina et al., 2010b; Jorjong et al., 2014; Dórea et al., 2017). The ROC curves plot the true positive rate or sensitivity versus the true negative rate (= 1 – specificity) for different thresholds applied on the predicted blood NEFA concentration. Only the most robust model, the one that performed best according to the procedure described in the previous section, was subjected to this ROC analysis. A separate analysis was done for the morning and evening observations of the validation set. The R package *pROC* version 1.13.0 (Robin et al., 2011) was used to calculate the ROC curves, to apply binormal smoothing to the ROC curves, to calculate the 95% confidence intervals (**CI**) of sensitivities, specificities and area under the curve (**AUC**) of the smoothed ROC curves and to statistically compare the smoothed ROC curves. The CI were calculated with 100 000 bootstrap replicates to obtain a fair estimate of the second significant digit (Fawcett, 2006). Statistical two-sided pairwise comparisons (α = 0.05) between ROC were done based on the area under the curve (AUC) and based on the sensitivities at given specificities from 0 to 1 in steps of 0.01, both using the bootstrap method with 100,000 replicates.

## RESULTS

### Data Exploration

The MIR transmittance spectra of the 1585 milk samples included in this study are presented in the top part of Figure 2 as the black solid and dotted lines. Lipid absorption peaks can clearly be observed as dips in the MIR transmittance spectra around 2928 & 2858 cm^−1^, 1745 cm^−1^, 1455 cm^−1^ and 1157 & 1078 cm^−1^, corresponding to respectively the C-H (alkyl) stretch, C=O (carbonyl) stretch, C-H bend and C-O stretch vibrations (Fox and McSweeney, 2006; De Marchi et al., 2009).

**Figure 2.**
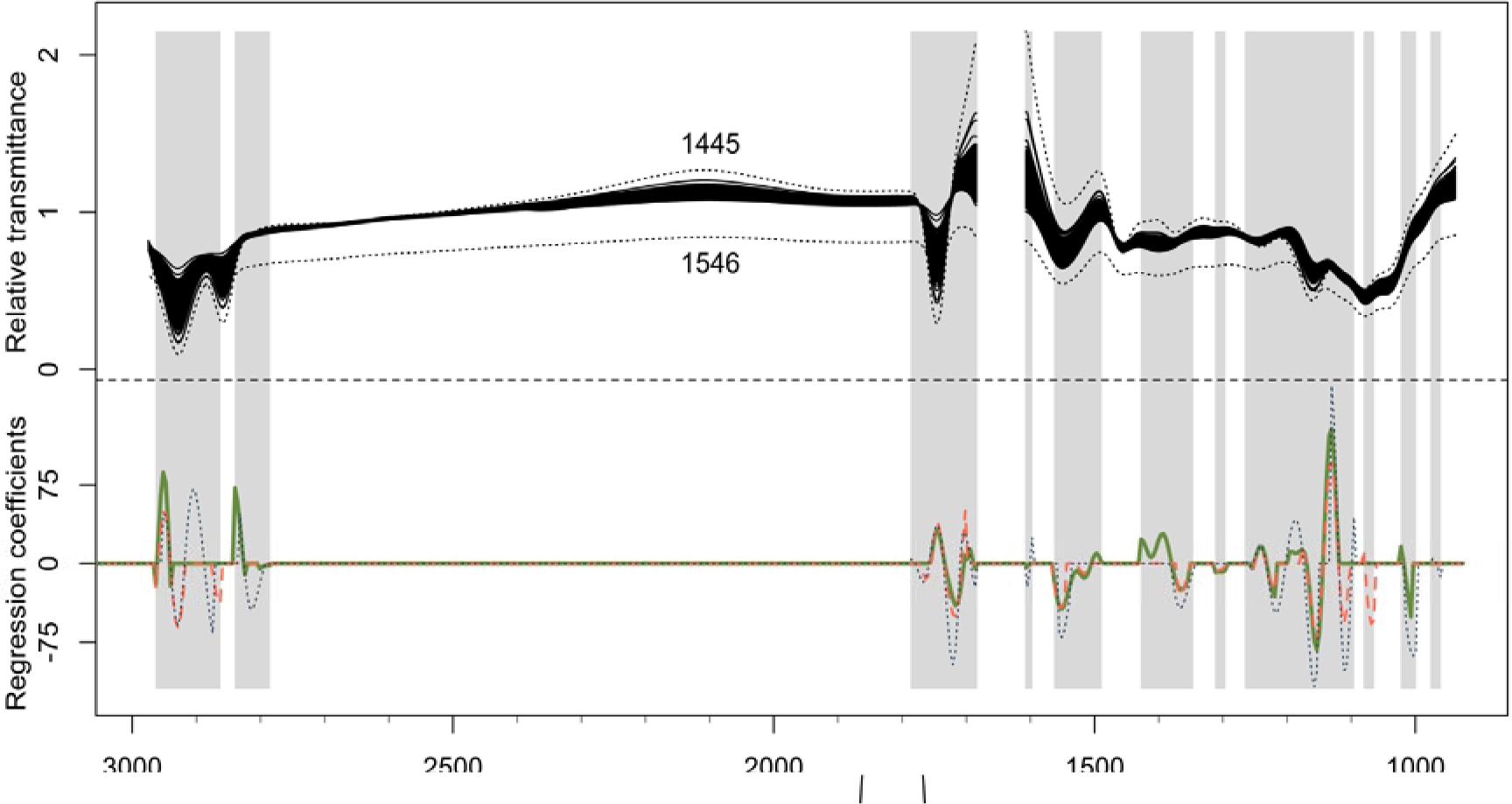
Top: Relative mid-infrared (MIR) transmittance spectra of the milk samples. The dotted black lines (with sample number) indicate 2 potential outlier spectra. The grey regions indicate the wavenumbers included in at least 1 of the 3 final partial least squares regression (PLSR) models (full, morning or evening) to predict the blood plasma non-esterified fatty acid concentration after applying a variable selection technique. Bottom: Regression coefficients for the 3 different PLSR models constructed using a calibration set with MIR spectra of respectively i) morning and evening milk samples (= full model, green solid), ii) only morning milk samples (= morning model, red dashed) and iii) only evening milk samples (= evening model, blue dotted) of herd A.

The Hotelling’s *T*^2^ and *Q*-statistics of the PCA model with 7 selected principal components and the scores for the first 2 principal components of that model were far beyond the 99% confidence limits for the spectra of sample 1445 (herd B, evening milking) and sample 1546 (herd C, morning milking), as shown in Appendix A1. Also, the raw transmittance spectra of these 2 outliers, illustrated with black dotted lines in top part of Figure 2, are clearly different from the other 1583 spectra, while the corresponding blood NEFA concentrations are not outlying. This suggests that these 2 samples have erroneous spectral measurements and were therefore removed from the dataset.

The reliability, accuracy, and robustness of spectroscopic calibrations are restricted to the range of the constituent of interest and the variation in measurement conditions taken into account during the calibration (Williams and Norris, 2001). The descriptive statistics of the blood NEFA concentrations linked to different subsets of milk MIR transmittance spectra are presented in Table 2. The entire calibration and validation set contain respectively 790 and 793 observations and they have a very similar mean, standard deviation and range for the blood NEFA. This table also illustrates the larger variability and range of the blood NEFA levels in herd A compared to herd B and C. Likely, this is the result of the higher number of blood samples being collected and analyzed (*n* = 658) and cows being monitored (*n* = 103) in herd A. Additionally, this might also be caused by differences in the genetic background and the management between the herds. The descriptive statistics for morning and evening samples in a same herd(s) are similar, but not exactly the same. This is because for some of the blood plasma samples only the respective morning or evening milk MIR spectra were collected and not both (Table 1).

**Table 2.**
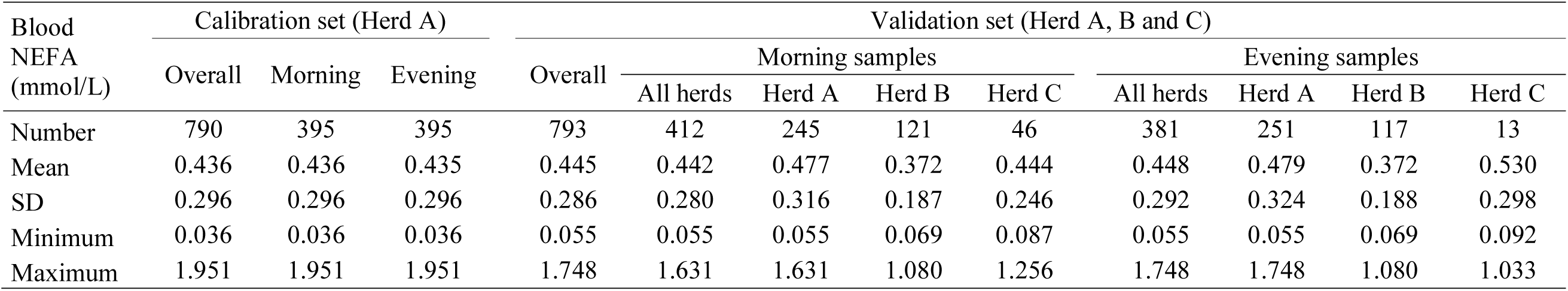
The descriptive statistics of the blood plasma non-esterified fatty acid (NEFA) concentrations linked to different subsets of milk mid-infrared transmittance spectra.

### Calibration on MIR Spectra of Morning and Evening Milk Samples (Full Model)

The 790 morning and evening observations of the calibration set were used to build a PLSR model (= full model) that relates the blood NEFA concentrations to the MIR transmittance spectra. The best performance was obtained when the MIR transmittance spectra were pre-processed using a 2^nd^ order Savitzky-Golay derivative with a window length of 7 wavenumber variables, followed by mean-centering. After this pre-processing step, the reversed interval PLSR method selected 117 wavenumbers that were most informative and resulted in the best model. The most informative regions of the MIR spectra are indicated as the grey regions in Figure 2.

Figure 3a shows the RMSECV and the RMSEP as a function of the number of latent variables included in the full PLSR model after applying the best pre-processing and selecting the best wavenumbers. A separate RMSEP is provided for the morning and evening observations of the validation set, respectively indicated with RMSEP_M_ and RMSEP_E_. The PLSR model with 6 latent variables, indicated with the green triangle, was finally selected. This model complexity resulted in nearly the minimum RMSEP_M_ and RMSEP_E_, confirming the right choice of number of latent variables based on the cross-validation and illustrating the robustness of the full model. Figure 3a clearly shows that the RMSEP_E_ is smaller than the RMSEP_M_ and that the latter is smaller than the RMSECV.

**Figure 3.**
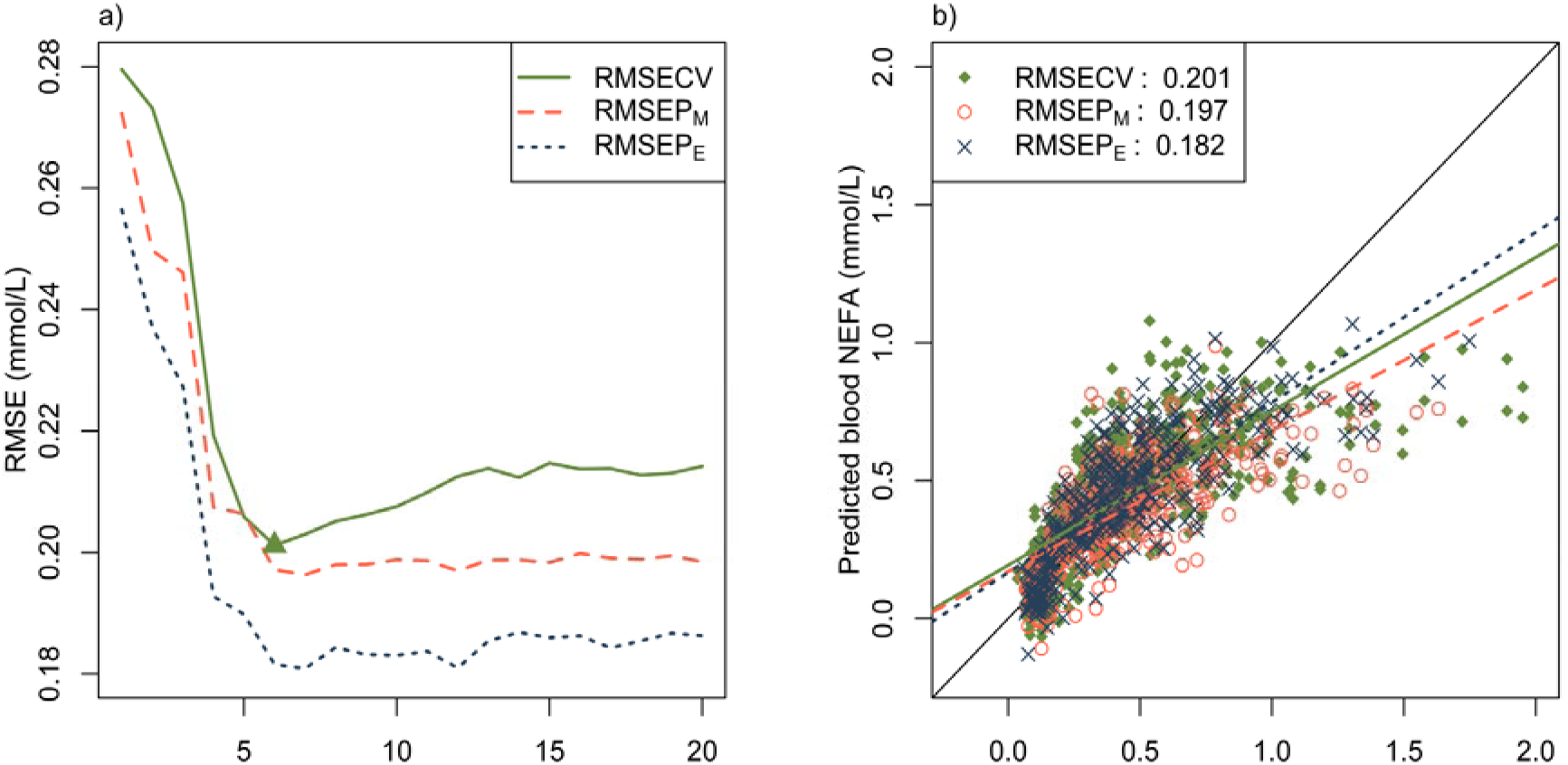
The results of the partial least squares regression (PLSR) model trained on a calibration set of mid-infrared transmittance spectra of milk samples collected during morning and evening milking sessions (= full model) on herd A to predict the non-esterified fatty acid (NEFA) concentration in the blood plasma of the respective cows for which blood was sampled in the morning. a) Root mean squared error (RMSE) for the calibration set in cross-validation (CV) and the morning (P_M_) and evening (P_E_) observations of the validation set (all 3 herds), in relation to the number of latent variables of the PLSR model. The green triangle indicates the number of selected latent variables (*n* = 6) for the final PLSR model. b) The predicted versus measured scatterplot for the calibration set (herd A) in cross-validation (CV) and the morning (P_M_) and evening (P_E_) observations of the validation set (3 herds).

The regression coefficients for the full model with 6 latent variables are presented with a green solid line in the bottom part of Figure 2. The regression coefficients follow a relatively smooth curve in function of the wavenumbers, which indicates that the PLSR model is not overfitting the calibration data. High absolute values for the regression coefficients were obtained around 2950 cm^−1^, 1750 cm^−1^ and 1150 – 990 cm^−1^, corresponding to important fat absorption bands: respectively the fat B, fat A and C-O stretch vibrations (Afseth et al., 2010). As the PLSR model uses the 2^nd^ derivative of the MIR spectra, some of the peaks in the regression coefficients are located at the flanks rather than the center of typical absorption peaks.

Figure 3b presents the predicted versus measured scatter plot for the full model with 6 latent variables. This figure illustrates that the prediction error of the full model varies a lot with the predicted blood NEFA concentration (y-axis), both for the cross-validated observations of the calibration set as well as for the morning and evening observations of the independent validation set. Additionally, the blood NEFA concentration is generally overestimated for true values (x-axis) between 0.2 and 0.55 mmol/L, while it is always underestimated for true concentrations above 1.2 mmol/L. The latter could explain why the RMSEP_M_ and RMSEP_E_ are lower than the RMSECV, as the validation set contains less observations with a very high blood NEFA concentration (Table 2). In Figure 3b, the predictions based on the evening observations of the validation set (blue crosses) are closer to the identity line compared to the ones based on the morning observations of the same set (red circles). This is the reason why the RMSEP_E_ values are smaller than the RMSEP_M_ values in Figure 3a and it suggests that the blood NEFA concentration in the morning can be predicted more accurately using MIR spectra of milk samples taken in the evening of that day rather than morning milk samples.

The prediction errors of the full model for different subsets of the validation set are summarized in Table 3. The heteroscedastic prediction error of the full model is clearly shown by the increasing RMSEP with increasing blood NEFA range (different horizontal sections of Table 3). Moreover, for the observations in the low blood NEFA range (0 – 0.6 mmol/L), the RMSEP values of the full model are all between 0.062 and 0.143 mmol/L, while for the middle blood NEFA range (0.6 – 1.2 mmol/L), the RMSEP values vary between 0.198 and 0.290 mmol/L. For the high blood NEFA range (1.2 – 2 mmol/L), the RMSEP values of the full model are between 0.620 and 0.793 mmol/L. Within the low, middle and high blood NEFA range, the RMSEP values do not differ much between herds, illustrating that the model can be used for new herds as well (cfr. herd B and C). Compared to the morning observations of the validation set, the RMSEP values for evening observations are in most cases slightly higher for the low blood NEFA range, while they are clearly lower for the complete, middle and high blood NEFA ranges. The observations based on the RMSEP values described in this paragraph are similar to the ones based on the RMSECV values obtained from the cross-validation of the calibration samples (results not shown), confirming the robustness of the full model.

**Table 3.** Root mean squared error of prediction (RMSEP) for the non-esterified fatty acid (NEFA) concentration in the blood plasma by the partial least squares regression models trained on morning and evening observations (= full model), only morning observations (= morning model) and only evening observations (= evening model) of the calibration set. The RMSEP values are provided for the different subsets (blood NEFA range, milking time and herd) of the validation set. Within each column and a specified blood NEFA concentration range, the RMSEP values with different subscripts indicate significant (α = 0.05) differences between the models according to Tukey HSD multiple comparison, with a letter lower in the alphabetical order indicating a better model.

| Blood NEFA<br>range<br>(mmol/L) | Model | Validation set - RMSEP (mmol/L) |  |  |  |  |  |  |  |  |
| --- | --- | --- | --- | --- | --- | --- | --- | --- | --- | --- |
|  |  | Overall | Morning observations |  |  |  | Evening observations |  |  |  |
|  |  |  | All herds | Herd A | Herd B | Herd C | All herds | Herd A | Herd B | Herd C |
| Complete:<br>0 – 2 | Full | 0.190<sup>a</sup> | 0.197<sup>a</sup> | 0.220<sup>a</sup> | 0.135 | 0.204 | 0.182<sup>a</sup> | 0.199<sup>a</sup> | 0.140<sup>a</sup> | 0.167 |
|  | Morning | 0.194<sup>b</sup> | 0.194<sup>a</sup> | 0.211<sup>a</sup> | 0.143 | 0.209 | 0.194<sup>b</sup> | 0.207<sup>b</sup> | 0.169<sup>b</sup> | 0.139 |
|  | Evening | 0.206<sup>b</sup> | 0.225<sup>b</sup> | 0.252<sup>b</sup> | 0.161 | 0.220 | 0.183<sup>a</sup> | 0.195<sup>a</sup> | 0.156<sup>a,b</sup> | 0.162 |
| Low:<br>0 – 0.6 | Full | 0.129<sup>a</sup> | 0.124<sup>a</sup> | 0.132 | 0.109 | 0.128 | 0.134<sup>a</sup> | 0.143<sup>b</sup> | 0.122<sup>a</sup> | 0.062 |
|  | Morning | 0.158<sup>b</sup> | 0.141<sup>b</sup> | 0.145 | 0.129 | 0.152 | 0.174<sup>b</sup> | 0.184<sup>c</sup> | 0.161<sup>b</sup> | 0.071 |
|  | Evening | 0.132<sup>a</sup> | 0.136<sup>a,b</sup> | 0.135 | 0.135 | 0.145 | 0.128<sup>a</sup> | 0.120<sup>a</sup> | 0.143<sup>a</sup> | 0.098 |
| Middle:<br>0.6 – 1.2 | Full | 0.241<sup>b</sup> | 0.269<sup>b</sup> | 0.270<sup>b</sup> | 0.253<sup>a,b</sup> | 0.290<sup>a,b</sup> | 0.208<sup>a,b</sup> | 0.198<sup>a,b</sup> | 0.230 | 0.258 |
|  | Morning | 0.211<sup>a</sup> | 0.234<sup>a</sup> | 0.232<sup>a</sup> | 0.218<sup>a</sup> | 0.277<sup>a</sup> | 0.185<sup>a</sup> | 0.174<sup>a</sup> | 0.219 | 0.206 |
|  | Evening | 0.281<sup>c</sup> | 0.328<sup>c</sup> | 0.339<sup>c</sup> | 0.286<sup>b</sup> | 0.322<sup>b</sup> | 0.223<sup>b</sup> | 0.221<sup>b</sup> | 0.226 | 0.231 |
| High:<br>1.2 – 2 | Full | 0.668<sup>b</sup> | 0.713<sup>b</sup> | 0.704<sup>b</sup> |  | 0.793 | 0.620<sup>b</sup> | 0.620<sup>b</sup> |  |  |
|  | Morning | 0.607<sup>a</sup> | 0.675<sup>a</sup> | 0.668<sup>a</sup> |  | 0.738 | 0.530<sup>a</sup> | 0.530<sup>a</sup> |  |  |
|  | Evening | 0.711<sup>c</sup> | 0.781<sup>c</sup> | 0.780<sup>c</sup> |  | 0.784 | 0.633<sup>b</sup> | 0.633<sup>b</sup> |  |  |

### Calibration on MIR Spectra of Morning or Evening Milk Samples

The PLSR model trained on the morning observations of the calibration set (= morning model), as well as the one trained on the evening observations (= evening model) provided the best results after applying a 2^nd^ order Savitzky-Golay derivative on the MIR transmittance spectra, followed by mean-centering. The optimal window length for the derivative was respectively 7 and 13 wavenumber variables. For both models, reversed interval PLSR proved to be the best variable selection method, resulting in respectively 92 and 126 retained wavenumbers (grey regions in Figure 2). Finally, 3 and 5 latent variables were selected for respectively the morning and evening PLSR model. The regression coefficients for the final morning and evening model are presented as respectively the red dashed and blue dotted lines in the bottom part of Figure 2. Both models have high absolute values for the regression coefficients in the regions near important MIR fat absorption bands, similar to the regression coefficients of the full model.

Analogues to the RMSEP values of the full model, the prediction errors of the morning and evening model for different subsets of the validation set are also provided in Table 3. The prediction performances of the three models (full, morning and evening) are compared for each subset of the validation set. Within each column (herd × milking time) and a specified blood NEFA range (complete, low, middle or high), RMSEP values with different subscripts indicate significant (α = 0.05) differences between the 3 models. Most subsets of herd C and some subsets involving herd B indicated no significant difference between the models. For those subsets, it was found that the statistical tests lacked power (β > 0.4) because of a too low number of samples. Therefore, the further discussion of the model comparison was only based on the tests with sufficient power (β < 0.2), which all happened to indicate a statistical effect of the model. The first column of Table 3 presents the RMSEP values for all the observations of the validation set in each of the 4 specified blood NEFA ranges.

In the complete range, the full model performs significantly better than the morning and the evening model, while there is no significant difference between the morning and the evening model. The full and evening models are not significantly different for the observations of the validation set in the low blood NEFA range, while they are both significantly better compared to the morning model. On the other hand, for the observations in the middle and high range, the morning model is significantly better than the full model, while the latter is better than the evening model.

For the low, middle and high blood NEFA range, the same trends are reflected in the different subsets of the validation set where the observations are split up per herd and/or milking time (Table 3, columns 2 to 9). For the complete blood NEFA range, the full model is significantly better compared to the evening model for all validation subsets with only morning observations, while it is significantly better compared to the morning model for all the subsets with evening observations. Additionally, it was found that the blood NEFA concentrations predicted with the morning model were on average 0.042 mmol/L higher compared to the predictions by the full model applied on the same milk MIR spectra, while the evening model resulted in blood NEFA predictions which were on average 0.048 mmol/L lower compared to the predictions by the full model. Given the fact that low blood NEFA concentrations are generally overestimated by the models, while the high blood NEFA concentrations are underestimated (Figure 3b), the morning model results in lower predictions in the high NEFA range, while the evening model results in lower prediction errors for the low NEFA range. This is also clearly reflected by the models’ RMSEP values for the different blood NEFA ranges. As the models mainly rely on the absorption by fat-related covalent bonds (Figure 2), the offset between the models probably results from the difference in average fat content between the morning milk samples (4.3%) and the evening milk samples (5.1%) involved in the training. Taken all this into account, it was concluded that the full model is the most robust of the 3 models and is therefore further explored in the ROC analysis in the next section.

Apart from the comparisons between the full, the morning and the evening model for each of the subsets of the validation set, a single 4-way ANOVA analysis was performed on the residuals of the observations in these different subsets. As all but one of the two-way interactions between the ANOVA factors were significant, the effect of the individual factors could not interpret independently from the other factors involved in the interaction(s). Accordingly, all combinations of the factors involved in these interactions were compared bilateral using a Tukey HSD multiple comparison. This analysis mainly points out that the prediction errors for the middle and high blood NEFA range are significantly higher compared to the complete and low range, but that the absolute levels of these errors depend on the model, the farm and the milking time.

Table 3 can also be used to study the difference in prediction error when 1 of the 3 models is applied on the different herds, or on either morning or evening observations. The RMSEP values for the 3 herds, except for the evening observations of herd C, are very close to each other for the same blood NEFA range (low, middle or high) for either morning or evening observations. This illustrates that the models can be easily transferred to new herds. The RMSEP values for the subsets of evening observations in herd C should be interpreted with caution as each of them is based on a low number of observations (*n* ≤ 13, Table 1). Comparing the prediction errors between morning and evening observations for respective subsets shows that the prediction errors for the morning observations are generally higher, especially for blood NEFA concentrations above 0.6 mmol/L (middle and high ranges). For the full model, a one-sided paired *T*-test applied on all the observations of the validation set pointed out that the blood NEFA predictions are more accurate (*p* = 0.009) if the model is applied on evening milk MIR spectra. This confirms that the blood NEFA concentration in the morning is predicted more accurately from milk MIR spectra taken during the evening milk session of the same day.

The observations based on the RMSEP values described in this section are similar to the ones based on the RMSECV values obtained from the cross-validation of the calibration samples (results not shown), confirming the robustness of the models and the validity of this analysis.

### Receiver Operating Characteristic Analysis of the Full Model

The smoothed ROC curves for the identification of detrimental blood NEFA concentrations based on the predictions of the full model are shown in Figure 4. Separate smoothed ROC curves are provided for the morning (red) and the evening observations (blue) of the validation set. The AUC of the smoothed ROC curves for the morning and evening observations are respectively 0.860 (95% CI: 0.815 – 0.901) and 0.898 (95% CI: 0.860 – 0.930). Accordingly, the AUC for the morning observations is significantly lower (p < 0.001) compared to the evening observations. Moreover, compared to the evening observations, the sensitivities for the morning observations are significantly lower in the range of specificities from 0.48 to 0.97. The average sensitivities are 0.752 and 0.573 for the morning observations and 0.831 and 0.690 for the evening observations (Figure 4) at specificities of respectively 0.8 and 0.9. Thus, cows with a detrimental blood NEFA concentration, as determined from their morning blood samples, can be detected more accurately using the MIR spectra of their milk collected during the evening milking session of that day. Moreover, it can identify 83 out of 100 cows with detrimental blood NEFA concentrations, while 20 out of 100 healthy cows will be wrongly classified as being at risk. Appendix A2 provides the mean values of the sensitivities and the 95% CI of the sensitivities and specificities at given specificities from 0.7 to 0.95 (in steps of 0.05) for the morning and the evening observations of the validation set.

**Figure 4.**
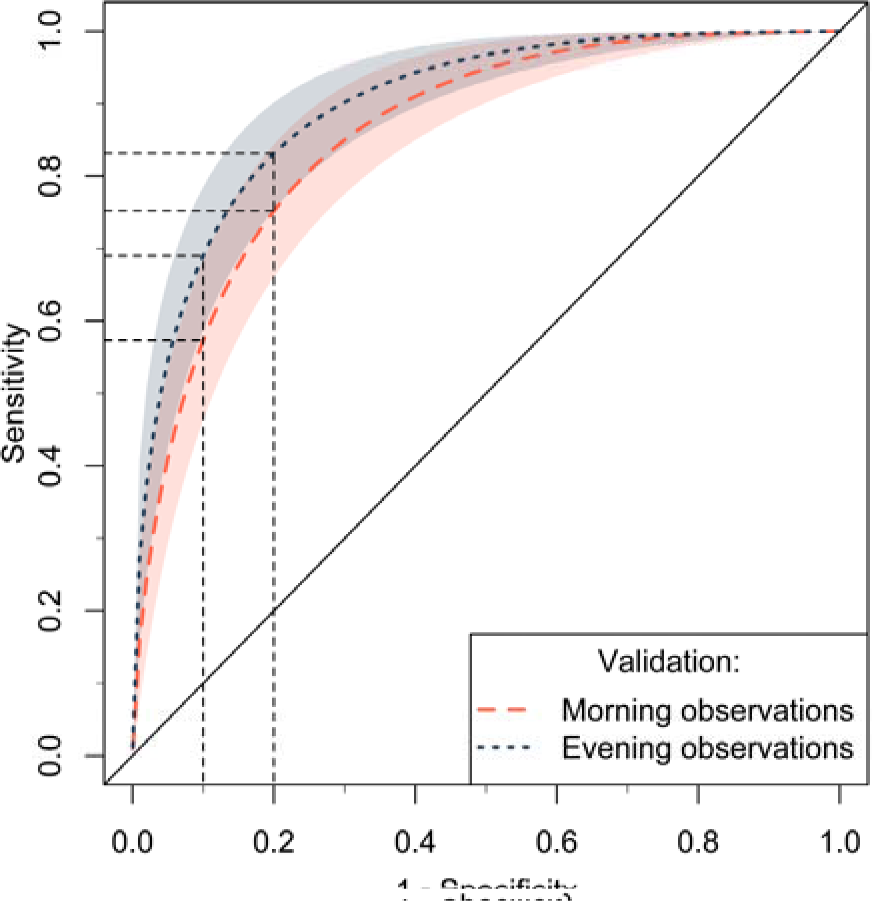
Smoothed receiver operating characteristic (ROC) curves for the identification of detrimental blood plasma non-esterified fatty acid (NEFA) concentrations (≥ 0.6 mmol/L). The NEFA concentrations were predicted with partial least squares regression models trained on morning and evening observations (= full model) of the calibration set. The mean values (lines) and 95% confidence intervals (areas) for the smoothed ROC curves are provided for morning (red, dashed) and evening observations (blue, dotted) of the validation set.

## DISCUSSION

### Morning and Evening Milk Samples

The validation of the PLSR models clearly indicates that, compared to morning milk, the MIR spectra of evening milk support more accurate predictions of the NEFA levels in blood taken in the morning of that day (Table 3, Figure 3 and Figure 4). Several studies have shown that blood NEFA follows a diurnal pattern with elevated levels from about 06:00 to 10:00 h in the morning, associated with a reduced energy intake during the night (Blum et al., 2000; Meier et al., 2010; Quiroz-Rocha et al., 2010). For this reason, extra attention was paid to the consistent timing of the blood and milk sampling. As the morning milking session was around 06:30 h, the majority of the period of expected elevated blood NEFA concentrations was after the morning milking session, thus mainly overlapping with the period in which the evening milk was produced. This likely introduced a time delay between the moment of elevated blood NEFA levels and the moment at which a change in the concentration of related milk biomarkers could be noticed, which is even further delayed by the metabolic processes in the liver that transfer NEFA into milk precursors and constituents (e.g. lipoproteins and ketone bodies). this delay would explain why the blood NEFA concentrations were predicted more accurately from MIR spectra of evening milk compared to morning milk as the blood samples were taken within 1 hour after the morning milking session, right in the middle of the time window of expected elevated blood NEFA levels. This hypothesis is also supported by the slightly higher NEFA levels predicted based on MIR spectra of evening milk samples compared to those based on the paired morning milk samples, especially for cows with detrimental blood NEFA concentrations (Figure 3b). It might be interesting for future research to study these dynamics more in detail by measuring the blood NEFA level at a frequent interval and investigating the link with the MIR spectra of morning and evening milk samples on that day and the days after.

The fact that the morning blood NEFA concentration can be predicted more accurately when the full model is applied on MIR spectra of evening milk rather than morning milk suggests that the evening milk samples contain more information on the morning blood NEFA concentration and/or that the morning milk samples are more subject to interfering effects. Nevertheless, training the model on solely evening milk MIR spectra (evening model) did not improve the prediction performance compared to the full model, even not if only MIR spectra of evening milk samples are considered in the validation. Moreover, the performance of the evening model was worse if applied on MIR spectra of morning milk samples. This suggests that including morning milk MIR spectra in the calibration set makes the prediction model more robust for potential interfering parameters that vary independent of the cow’s blood NEFA level. One of these interfering effects might be the total fat content in the milk, which is generally higher in evening milk compared to morning milk (Forsbäck et al., 2010). Apart from the results obtained in our study, the full model is probably more robust under practical conditions where farms have varying milking and feeding frequencies.

### Prediction of Blood Plasma NEFA Concentration

The regression coefficients in Figure 2 show that the PLSR models primarily use information from the fat-related MIR absorption bands. During negative energy status, excessive amounts of NEFA are mobilized from the adipose tissue and part of them is transferred to the milk. These NEFA are particularly rich in long-chain fatty acids (**FA**), such as C18:1 FA (Jorjong et al., 2014). Dórea et al. (2017) found a nonlinear relation (*R*^2^ = 0.42 and *p* < 0.001) between the concentrations of NEFA in the blood plasma and C18:1 FA in the milk fat. Moreover, the milk C18:1 FA increased nearly linearly with increasing blood NEFA for blood NEFA levels below 400 µEq/L, while the milk C18:1 FA concentration was practically constant for blood NEFA concentrations above 800 µEq/L. This suggests that the C18:1 FA concentration in milk fat saturates when the blood NEFA increases above a certain concentration. On the other hand, Jorjong et al. (2014) suggested a linear relation (*R*^2^ = 0.383) between the concentrations of NEFA in the blood plasma and C18:1 *cis*-9 FA in the milk fat. However, their linear function slightly underestimated the milk C18:1 *cis*-9 FA for blood NEFA concentrations between 0.2 and 0.4 mmol/L, while it overestimated the milk C18:1 cis-9 FA for blood NEFA levels below 0.1 and above 0.9 mmol/L. Accordingly, the data of Jorjong et al. (2014) confirms the non-linear trend found by Dórea et al. (2017). In our study, the predicted blood NEFA concentrations versus the actual blood NEFA levels (Figure 3b) follows a very similar nonlinear trend as the milk C18:1 FA in the studies of Dórea et al. (2017) and Jorjong et al. (2014). Therefore, it is likely that our PLSR models largely rely on the MIR absorption by C18:1 and related FA in milk. Several researchers already explored MIR spectroscopy to predict the concentration of certain FA in milk, obtaining *R*^2^ values for the prediction of C18:1 FA between 0.11 and 0.96 (Rutten et al., 2009; Afseth et al., 2010; Soyeurt et al., 2011). Mäntysaari et al. (2019) used the PLSR models developed by Soyeurt et al. (2011) to predict the milk FA concentrations from milk MIR spectra and accordingly studied the relation between the predicted milk FA and the blood NEFA concentration. It was found that C18:1 *cis*-9 and the sum of C18:1 FA in milk had the highest correlation (*r* = 0.73) with blood NEFA, confirming our hypothesis.

The predicted versus measured scatterplot in Figure 3b, as well as the RMSEP values in Table 3, clearly show that the accuracy of the prediction of the blood NEFA from milk MIR spectra is limited, especially if the blood NEFA concentration is high. The full model results in RMSEP values of 0.197, 0.182 and 0.190 mmol/L when evaluated on respectively morning observations, evening observations or a mixed set of morning and evening observations of the validation set. Taking into account the standard deviations of the blood NEFA concentration for the different sets (Table 2), the *R*^2^ values are respectively 0.502, 0.613 and 0.558. Nevertheless, the RMSEP and *R*^2^ values strongly depend, because of heteroscedasticity, on the proportion of observations with a high blood NEFA concentration in the respective datasets. To account for this non-linear effect, we also explored non-linear models, such as convolutional neural networks, and a logarithmic transformation of the blood NEFA levels before applying PLSR without success. Moreover, using a more balanced calibration set with a similar number of observations with high and low blood NEFA levels through bootstrapping did not improve the performance of the prediction model either (results not shown). Because of the heteroscedasticity of the prediction error, benchmarking our results against earlier studies is challenging and should be done with caution.

Dórea et al. (2017) obtained RMSE values of 169 – 220 µEq/L (equivalent to µmol/L) and *R*^2^ values of 0.080 – 0.457 for the prediction of blood NEFA levels for individual cows from different linear combinations or ratios of milk FA concentrations obtained from GLS analysis. As the descriptive statistics for the blood NEFA are very similar in their dataset and ours (Table 2), it is fair to compare the results of these 2 studies. The prediction errors reported by Dórea et al. are very close to the ones obtained in our study. Nevertheless, the performances of their models are only reported for the calibration set and thus might be overoptimistic. Additionally, the approach followed by Dórea et al. requires labor and cost intensive FA isolation and GLS analysis.

Mäntysaari et al. (2019) used a linear combination of C18:1 *cis*-9 and medium chain FA concentrations in milk, derived from evening milk MIR spectra, and lactation stage to predict the morning blood NEFA concentration, obtaining an *R*^2^ of 0.61 and an RMSECV of 0.182 mmol/L. A similar approach using morning milk MIR spectra resulted in an *R*^2^ of 0.52 and an RMSECV of 0.198 mmol/L. Although these results only represent the cross-validation of the model, and thus might be overoptimistic, they are in close agreement with the results obtained for the independent validation in our study.

Recently, Benedet et al. (2019), Grelet et al. (2019) and Luke et al. (2019) published PLSR models that predict the blood NEFA levels directly from the MIR spectra of raw milk samples. The prediction performance of Grelet’s model (*R*^2^ = 0.39 and RMSECV = 344 µeq/L) is only based on cross-validation of the calibration set and should thus be confirmed on an external validation set (Grelet et al., 2019). Still, their results are inferior to the ones obtained in our study due to the higher prediction error by Grelet’s model in the low blood NEFA range. Moreover, while our full model is relatively accurate (RMSEP ≤ 0.143 mmol/L) in this range, Grelet’s model generally overestimates the low blood NEFA concentrations. In the high blood NEFA range, Grelet’s model performs similar to our models, both underestimating the blood NEFA concentration. As a result, the prediction error of Grelet’s model is nearly homoscedastic, but worse compared to our model, especially in the low blood NEFA range. Benedet et al. (2019) obtained a PLSR model that performed better, compared to Grelet’s model, with an *R*^2^ of 0.52 and a standard error of prediction 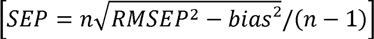 of 0.24 mmol/L for a randomly selected validation set. Still, these results are slightly worse compared to the ones obtained in our study. In contrast to our models, Benedet’s model only uses wavenumbers between 1450 and 1000 cm^−1^ and thus ignores the fat absorption bands at around 2928, 2858 and 1745 cm^−1^. Additionally, it should be taken into account that the model performance typically deteriorates when it is applied on a completely independent herd, as illustrated by Luke et al. (2019). Moreover, the *R*^2^ of Luke’s model dropped from 0.61 for a randomly selected validation set, covering the same herds as the calibration set, to 0.45 for a totally independent herd. The better performance of our full model is likely the result of a higher number of calibration samples (*n* = 790) in combination with a well-controlled timing and protocol for blood and milk sample collection (Quiroz-Rocha et al., 2010).

### Receiver Operating Characteristic Analysis

Unlike the regression analysis, in which the RMSE is subject to the effect of heteroscedasticity, the result of the ROC analysis is less dependent on the relative number of observations with a high blood NEFA concentration. Accordingly, these ROC analyses allow for a more objective comparison among different studies. Dórea et al. (2017) obtained their best results to identify cows with detrimental blood NEFA concentrations (≥ 600 µEq/L) based on the milk C13:0 FA using a threshold of 0.036 g FA per 100 g milk fat. This resulted in an AUC, sensitivity and specificity of respectively 0.90, 0.859 and 0.823. The ROC curve to detect detrimental blood NEFA levels based on milk C18:1 *cis*-9 FA and reported by Jorjong et al. (2014) had a sensitivity of 0.75 and 0.5 at a specificity of respectively 0.79 and 0.935. The full model obtained in our approach and applied on the evening observations results in an AUC of 0.898 and a sensitivity of 0.831 at a specificity of 0.8. The same model applied on morning observations has an AUC of 0.860 and a sensitivity of 0.752 at a specificity of 0.8 (Appendix A2). Therefore, it can be concluded that our model, which only requires MIR spectral analysis of raw milk, is not inferior compared to more complex techniques that require characterization of certain FA in the milk fat. A similar approach followed by Luke et al. (2019) to identify elevated blood NEFA levels resulted in a AUC values of 0.87 and 0.82, sensitivities of 0.73 and 0.25 and specificities of 0.81 and 0.90 for respectively a randomly selected and a completely independent validation set. Thus, our model tends to be slight more robust compared to the one of Luke et al. (2019).

Although the prediction accuracy is not excellent, the developed model can provide valuable information to further improve genetics, nutrition and management of dairy cows. As it can be applied on millions of historical and future milk MIR spectra, this approach can reveal detailed information on the energy status of individual cows, herds and pedigrees. This could potentially result in improved estimations of breeding values and the identification of specific genetic markers for metabolic resilience.

## CONCLUSIONS

In this study, we successfully predicted the blood NEFA level in individual dairy cows from their milk MIR spectra. The best model was obtained after training on MIR spectra of both morning and evening milk samples. The NEFA concentration of blood plasma samples taken in the morning were predicted with a higher accuracy if the model was applied on MIR spectra of evening milk samples. The obtained prediction accuracy is acceptable for low blood NEFA levels, but is unsatisfactory if the blood NEFA concentration is high. Nevertheless, low and intermediate/high blood NEFA levels could be discriminated well to identify 83 out of 100 cows with detrimental blood NEFA levels, while only 20 out of 100 healthy cows are wrongly classified. This opens opportunities for identifying cows at risk of a negative energy status and studying the metabolic resilience of individual cows and pedigrees.

## ACKNOWLEDGEMENTS

This study was funded by the Finnish Ministry of Agriculture and Forestry (1844/312/2012), Valio Ltd, Faba co-op, Viking Genetics, Finnish Cattle Breeding Foundation and Raisioagro Ltd. Ben Aernouts and Ines Adriaens were respectively funded as postdoctoral fellow (11ZG916N) and aspirant fellow (12K3916N) by the Research Foundation Flanders (FWO, Brussels, Belgium). Ben Aernouts obtained additional funding from the Research Foundation Flanders to perform a long research stay abroad at the Natural Resources Institute of Finland (Luke), grant V407018N.

## APPENDICES

**Appendix A1.**
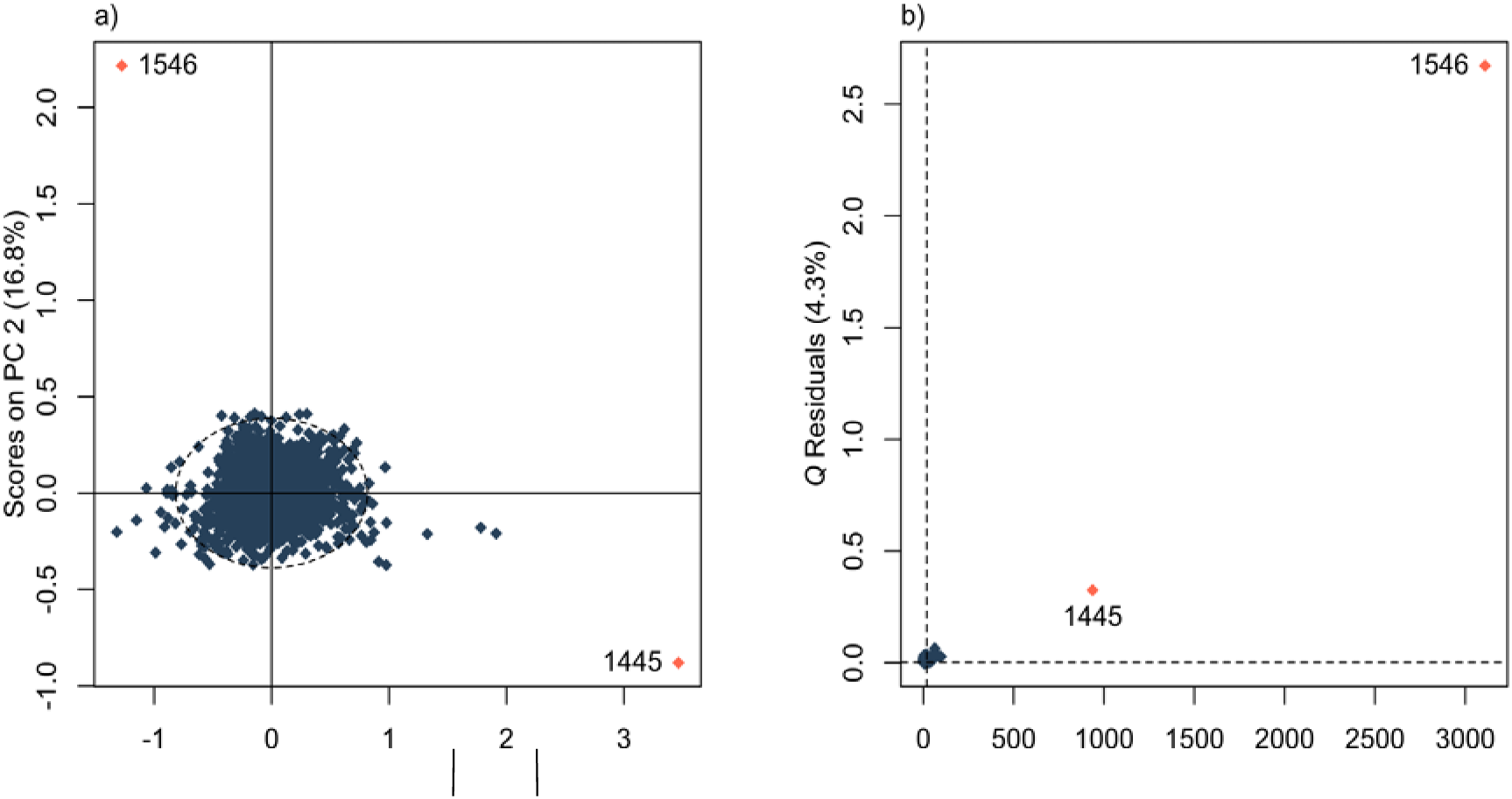
Figures of a) The scores plot of principal component (PC) 1 versus PC2 of the principal component analysis (PCA) on all 1585 mean-centered mid-infrared transmittance spectra. The dashed ellipse represents the 99% confidence limits of the scores on PC1 and PC2. b) The influence plot (*Q* residual versus Hotelling *T*^2^ statistics) for the PCA model with 7 PC presenting the of the PCA. The dashed lines represent the 99% confidence limits on respectively the 2 statistics. In both figures, each dot represents a different sample spectrum and the red dots (with sample number) indicate potential outlier spectra.

**Appendix A2.**
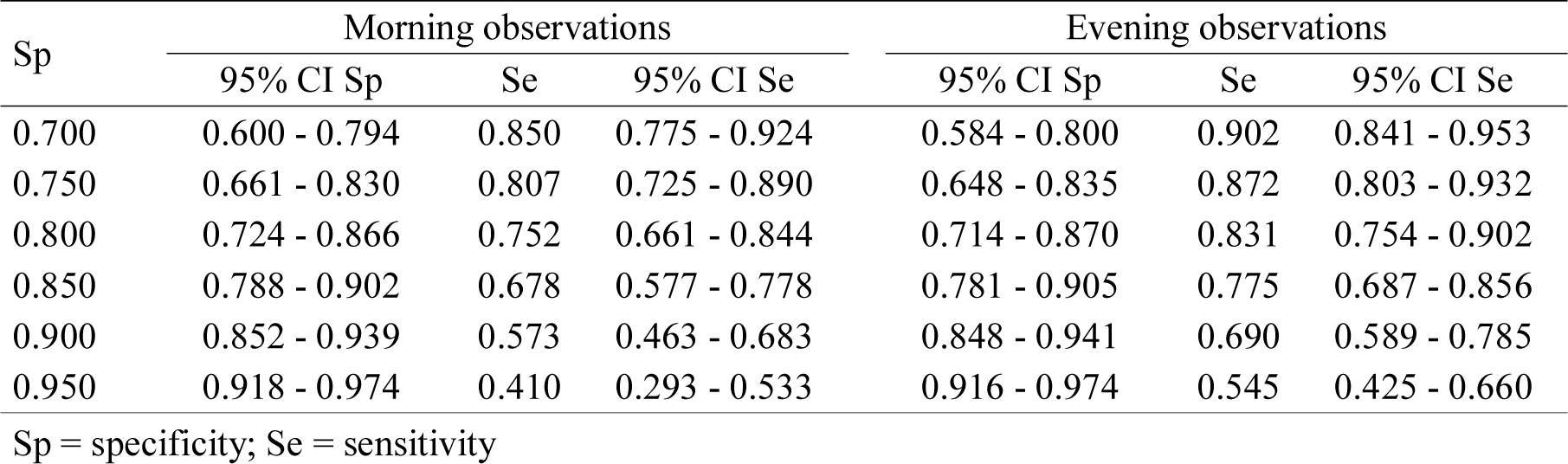
Table with the mean values of the sensitivities and the 95% confidence intervals (CI) of the sensitivities and specificities at given specificities for the morning and the evening observations of the validation set.

| Sp | Morning observations |  |  | Evening observations |  |  |
| --- | --- | --- | --- | --- | --- | --- |
|  | 95% CI Sp | Se | 95% CI Se | 95% CI Sp | Se | 95% CI Se |
| 0.700 | 0.600 - 0.794 | 0.850 | 0.775 - 0.924 | 0.584 - 0.800 | 0.902 | 0.841 - 0.953 |
| 0.750 | 0.661 - 0.830 | 0.807 | 0.725 - 0.890 | 0.648 - 0.835 | 0.872 | 0.803 - 0.932 |
| 0.800 | 0.724 - 0.866 | 0.752 | 0.661 - 0.844 | 0.714 - 0.870 | 0.831 | 0.754 - 0.902 |
| 0.850 | 0.788 - 0.902 | 0.678 | 0.577 - 0.778 | 0.781 - 0.905 | 0.775 | 0.687 - 0.856 |
| 0.900 | 0.852 - 0.939 | 0.573 | 0.463 - 0.683 | 0.848 - 0.941 | 0.690 | 0.589 - 0.785 |
| 0.950 | 0.918 - 0.974 | 0.410 | 0.293 - 0.533 | 0.916 - 0.974 | 0.545 | 0.425 - 0.660 |

Sp = specificity; Se = sensitivity

## REFERENCES

Adewuyi, A.A., E. Gruysi, and F.J.C.M.V. Eerdenburg. 2005. Non esterified fatty acids (NEFA) in dairy cattle. A review. Vet. Q. 27:117–126. doi:10.1080/01652176.2005.9695192.

Aernouts, B., E. Polshin, W. Saeys, and J. Lammertyn. 2011. Mid-infrared spectrometry of milk for dairy metabolomics: a comparison of two sampling techniques and effect of homogenization. Anal. Chim. Acta 705:88–97. doi:10.1016/j.aca.2011.04.018.

Afseth, N.K., H. Martens, Å. Randby, L. Gidskehaug, B. Narum, K. Jørgensen, S. Lien, and A. Kohler. 2010. Predicting the fatty acid composition of milk: A comparison of two fourier transform infrared sampling techniques. Appl. Spectrosc. 64:700–707. doi:10.1366/000370210791666200.

Barnes, R.J., M.S. Dhanoa, and S.J. Lister. 1989. Standard normal variate transformation and de-trending of near-infrared diffuse reflectance spectra. Appl. Spectrosc. 43:772–777. doi:10.1366/0003702894202201.

Beer. 1852. Bestimmung der Absorption des rothen Lichts in farbigen Flüssigkeiten. Ann. Phys. 162:78–88. doi:10.1002/andp.18521620505.

Benedet, A., M. Franzoi, M. Penasa, E. Pellattiero, and M. De Marchi. 2019. Prediction of blood metabolites from milk mid-infrared spectra in early-lactation cows. J. Dairy Sci.. doi:10.3168/jds.2019-16937.

Bjerre-Harpøth, V., N.C. Friggens, V.M. Thorup, T. Larsen, B.M. Damgaard, K.L. Ingvartsen, and K.M. Moyes. 2012. Metabolic and production profiles of dairy cows in response to decreased nutrient density to increase physiological imbalance at different stages of lactation. J. Dairy Sci. 95:2362–2380. doi:10.3168/jds.2011-4419.

Blum, J.W., R.M. Bruckmaier, P.Y. Vacher, A. Münger, and F. Jans. 2000. Twenty-Four-Hour Patterns of Hormones and Metabolites in Week 9 and 19 of Lactation in High-Yielding Dairy Cows fed Triglycerides and Free Fatty Acids. J. Vet. Med. Ser. A Physiol. Pathol. Clin. Med. 47:43–60. doi:10.1046/j.1439-0442.2000.00266.x.

Bro, R., and A.K. Smilde. 2014. Principal component analysis. Anal. Methods 6:2812–2831. doi:10.1039/c3ay41907j.

Cederkvist, H.R., A.H. Aastveit, and T. Næs. 2005. A comparison of methods for testing differences in predictive ability. J. Chemom. 19:500–509. doi:10.1002/cem.956.

Chong, I.G., and C.H. Jun. 2005. Performance of some variable selection methods when multicollinearity is present. Chemom. Intell. Lab. Syst. 78:103–112. doi:10.1016/j.chemolab.2004.12.011.

Dórea, J.R.R., E.A. French, and L.E. Armentano. 2017. Use of milk fatty acids to estimate plasma nonesterified fatty acid concentrations as an indicator of animal energy balance. J. Dairy Sci. 100:6164–6176. doi:10.3168/jds.2016-12466.

Enjalbert, F., M.C. Nicot, C. Bayourthe, and R. Moncoulon. 2010. Ketone Bodies in Milk and Blood of Dairy Cows: Relationship between Concentrations and Utilization for Detection of Subclinical Ketosis. J. Dairy Sci. 84:583–589. doi:10.3168/jds.s0022-0302(01)74511-0.

Fawcett, T. 2006. An Introduction to ROC analysis. Pattern Recognit. Lett. 27:861–874. doi:10.1016/j.patrec.2005.10.010.

Forsbäck, L., H. Lindmark-Månsson, A. Andrén, M. Akerstedt, L. Andrée, and K. Svennersten-Sjaunja. 2010. Day-to-day variation in milk yield and milk composition at the udder-quarter level. J. Dairy Sci. 93:3569–3577. doi:10.3168/jds.2009-3015.

Fox, P.F., and P.L.H. McSweeney. 2006. Advanced Dairy Chemistry Volume 2: Lipids. 3rd ed. Springer, New York, USA.

Franzoi, M., G. Niero, G. Visentin, M. Penasa, M. Cassandro, and M. De Marchi. 2019. Variation of Detailed Protein Composition of Cow Milk Predicted from a Large Database of Mid-Infrared Spectra. Animals 9:176. doi:10.3390/ani9040176.

Geladi, P., D. MacDougall, and H. Martens. 1985. Linearization and Scatter-Correction for Near-Infrared Reflectance Spectra of Meat. Appl. Spectrosc. 39:491–500. doi:10.1366/0003702854248656.

Grelet, C., C. Bastin, M. Gelé, J.-B. Davière, M. Johan, A. Werner, R. Reding, J.A. Fernandez Pierna, F.G. Colinet, P. Dardenne, N. Gengler, H. Soyeurt, and F. Dehareng. 2016. Development of Fourier transform mid-infrared calibrations to predict acetone, β-hydroxybutyrate, and citrate contents in bovine milk through a European dairy network. J. Dairy Sci. 99:4816–4825. doi:10.3168/jds.2015-10477.

Grelet, C., J.A. Fernández Pierna, P. Dardenne, V. Baeten, and F. Dehareng. 2015. Standardization of milk mid-infrared spectra from a European dairy network. J. Dairy Sci. 98:2150–2160. doi:10.3168/jds.2014-8764.

Grelet, C., A. Vanlierde, M. Hostens, L. Foldager, M. Salavati, K.L. Ingvartsen, M. Crowe, M.T. Sorensen, E. Froidmont, C.P. Ferris, C. Marchitelli, F. Becker, T. Larsen, F. Carter, and F. Dehareng. 2019. Potential of milk mid-IR spectra to predict metabolic status of cows through blood components and an innovative clustering approach. Animal 13:649–658. doi:10.1017/S1751731118001751.

Van Haelst, Y.N.T., A. Beeckman, A.T.M. Van Knegsel, and V. Fievez. 2008. Short Communication: Elevated Concentrations of Oleic Acid and Long-Chain Fatty Acids in Milk Fat of Multiparous Subclinical Ketotic Cows. J. Dairy Sci. 91:4683–4686. doi:10.3168/jds.2008-1375.

ICAR. 2019. Section 12 – Guidelines for Milk Analysis.

ISO. 2013. Milk and liquid milk products -- Guidelines for the application of mid-infrared spectrometry. Page 14 in International Standard ISO 9622:2013/IDF 141:2013. International Dairy Federation.

Jorjong, S., A.T.M. van Knegsel, J. Verwaeren, M.V. Lahoz, R.M. Bruckmaier, B. De Baets, B. Kemp, and V. Fievez. 2014. Milk fatty acids as possible biomarkers to early diagnose elevated concentrations of blood plasma nonesterified fatty acids in dairy cows. J. Dairy Sci. 97:7054–7064. doi:10.3168/jds.2014-8039.

Kemps, B.J., W. Saeys, K. Mertens, P. Darius, J.G. De Baerdemaeker, and B. De Ketelaere. 2010. The importance of choosing the right validation strategy in inverse modelling. J. Near Infrared Spectrosc. 18:231–237. doi:10.1255/jnirs.882.

Leblanc, S. 2010. Monitoring Metabolic Health of Dairy Cattle in the Transition Period. J. Reprod. Dev. 56:S29–S35. doi:10.1262/jrd.1056S29.

Leroy, J.L.M.R., P. Bossaert, G. Opsomer, and P.E.J. Bols. 2011. The effect of animal handling procedures on the blood non-esterified fatty acid and glucose concentrations of lactating dairy cows. Vet. J. 187:81–84. doi:10.1016/j.tvjl.2009.10.003.

Leroy, J.L.M.R., T. Vanholder, B. Mateusen, A. Christophe, G. Opsomer, A. de Kruif, G. Genicot, and A. Van Soom. 2005. Non-esterified fatty acids in follicular fluid of dairy cows and their effect on developmental capacity of bovine oocytes in vitro. Reproduction 130:485–495. doi:10.1530/rep.1.00735.

Luke, T.D.., S. Rochfort, W.J. Wales, V. Bonfatti, L. Marett, and J.E. Pryce. 2019. Metabolic profiling of early-lactation dairy cows using milk mid-infrared spectra. J. Dairy Sci. 102:1747–1760. doi:10.3168/jds.2018-15103.

Mäntysaari, P., E.A. Mäntysaari, T. Kokkonen, T. Mehtiö, S. Kajava, C. Grelet, P. Lidauer, and M.H. Lidauer. 2019. Body and milk traits as indicators of dairy cow energy status in early lactation. J. Dairy Sci. 102:7904–7916. doi:10.3168/jds.2018-15792.

De Marchi, M., C.C. Fagan, C.P. O’Donnell, A. Cecchinato, R. Dal Zotto, M. Cassandro, M. Penasa, and G. Bittante. 2009. Prediction of coagulation properties, titratable acidity, and pH of bovine milk using mid-infrared spectroscopy. J. Dairy Sci. 92:423–432. doi:10.3168/jds.2008-1163.

De Marchi, M., V. Toffanin, M. Cassandro, and M. Penasa. 2014. Invited review: Mid-infrared spectroscopy as phenotyping tool for milk traits1. J. Dairy Sci. 97:1171–1186. doi:10.3168/jds.2013-6799.

Martens, H., and T. Næs. 1987. Multivariate calibration by data compression. P.C. Williams and K. Norris, ed. American Association of Cereal Chemists, St Paul, MN, USA.

Meier, S., E.S. Kolver, G.A. Verkerk, and J.R. Roche. 2010. Effects of divergent Holstein-Friesian strain and diet on diurnal patterns of plasma metabolites and hormones. J. Dairy Res. 77:432–437. doi:10.1017/S002202991000052X.

Norgaard, L., J. Wagner, J.P. Nielsen, L. Munc, and S.B. Engelsen. 2000. Interval Partial Least-Squares Regression (iPLS): A comparative chemometric study with an example from Near-Infrared Spectroscopy. Appl. Spectrosc. 54:413–419. doi:10.1366/0003702001949500.

Ospina, P.A., D.V. Nydam, T. Stokol, and T.R. Overton. 2010a. Associations of elevated nonesterified fatty acids and β-hydroxybutyrate concentrations with early lactation reproductive performance and milk production in transition dairy cattle in the northeastern United States. J. Dairy Sci. 93:1596–1603. doi:10.3168/jds.2009-2852.

Ospina, P.A., D.V. Nydam, T. Stokol, and T.R. Overton. 2010b. Evaluation of nonesterified fatty acids and β-hydroxybutyrate in transition dairy cattle in the northeastern United States: Critical thresholds for prediction of clinical diseases. J. Dairy Sci. 93:546–554. doi:10.3168/jds.2009-2277.

Quiroz-Rocha, G.F., S.J. LeBlanc, T.F. Duffield, B. Jefferson, D. Wood, K.E. Leslie, and R.M. Jacobs. 2010. Short communication: Effect of sampling time relative to the first daily feeding on interpretation of serum fatty acid and β-hydroxybutyrate concentrations in dairy cattle. J. Dairy Sci. 93:2030–2033. doi:10.3168/jds.2009-2141.

R Core Team. 2017. R: A language and environment for statistical computing. Vienna, Austria.

Robin, X., N. Turck, A. Hainard, N. Tiberti, F. Lisacek, J. Sanchez, and M. Müller. 2011. pROC: an open-source package for R and S+ to analyze and compare ROC curves. BMC Bioinformatics 12:1–8.

Ruckstuhl, A.F., M.P. Jacobson, R.W. Field, and J.A. Dodd. 2001. Baseline subtraction using robust local regression estimation. J. Quant. Spectrosc. Radiat. Transf. 68:179–193. doi:10.1016/S0022-4073(00)00021-2.

Rutten, M.J.M., H. Bovenhuis, K.A. Hettinga, H.J.F. van Valenberg, and J.A.M. van Arendonk. 2009. Predicting bovine milk fat composition using infrared spectroscopy based on milk samples collected in winter and summer. J. Dairy Sci. 92:6202–6209. doi:10.3168/jds.2009-2456.

Salin, S., J. Taponen, K. Elo, I. Simpura, A. Vanhatalo, R. Boston, and T. Kokkonen. 2012. Effects of abomasal infusion of tallow or camelina oil on responses to glucose and insulin in dairy cows during late pregnancy. J. Dairy Sci. 95:3812–25. doi:10.3168/jds.2011-5206.

Savitzky, A., and M.J.E. Golay. 1964. Smoothing and Differentiation of Data by Simplified Least Squares Procedures. Anal. Chem. 36:1627–1639. doi:10.1021/ac60214a047.

Scalia, D., N. Lacetera, U. Bernabucci, K. Demeyere, L. Duchateau, and C. Burvenich. 2006. In Vitro Effects of Nonesterified Fatty Acids on Bovine Neutrophils Oxidative Burst and Viability. J. Dairy Sci. 89:147–154. doi:10.3168/jds.s0022-0302(06)72078-1.

Snee, R. 1977. Validation of regression models: methods and examples. Technometrics 19:415–428. doi:10.2307/1267881.

Soyeurt, H., D. Bruwier, J.-M. Romnee, N. Gengler, C. Bertozzi, D. Veselko, and P. Dardenne. 2009. Potential estimation of major mineral contents in cow milk using mid-infrared spectrometry. J. Dairy Sci. 92:2444–2454. doi:10.3168/jds.2008-1734.

Soyeurt, H., F. Dehareng, N. Gengler, S. Mcparland, E. Wall, D.P. Berry, and M. Coffey. 2011. Mid-infrared prediction of bovine milk fatty acids across multiple breeds, production systems, and countries. J. Dairy Sci. 94:1657–1667. doi:10.3168/jds.2010-3408.

Westad, F., and H. Martens. 2000. Variable selection in near infrared spectroscopy based on significance testing in partial least squares regression. J. Near Infrared Spectrosc. 8:117–124. doi:10.1255/jnirs.271.

Williams, P., and K. Norris. 2001. Near-Infrared Technology in the Agricultural and Food Industries. 2nd ed. American Association of Cereal Chemist, St. Paul, USA.

